# Consequences of life-cycle complexity to the potential for evolutionary branching

**DOI:** 10.1101/2022.08.31.506002

**Authors:** Paula Vasconcelos, Marco Saltini, Claus Rueffler

**Affiliations:** Department of Ecology and Genetics, Animal Ecology, Uppsala University, Norbyvägen 18D, 752 36 Uppsala, Sweden; Swedish Collegium for Advanced Study (SCAS), Thunbergsvägen 2, 752 38 Uppsala, Sweden

**Keywords:** Adaptive dynamics, coexistence, evolutionary branching, ontogentic niche shift, disruptive selection, resource competition, resource polymorphism

## Abstract

Complex life-cycles – that is, organismal development that unfolds across ecological niches – are pervasive in nature. In this work we set out to investigate the effects of complex life-cycles on the potential for diversification via evolutionary branching. We did this by analyzing a mathematical model of a consumer with two life-stages, each of which is characterized by a specific feeding efficiency trait that undergoes evolutionary change in response to ecological conditions such as resource competition. We find (i) that life-cycle complexity can favor diversification when compared to simple life-cycles, as there is a larger potential for evolutionary branching in the trait of the life-stage that has a higher population density; (ii) that evolution favors character displacement to minimize intra-stage resource competition; and (iii) that under certain parameters more than one evolutionary branching event can occur.

## Introduction

The potential effects of complex life-cycles on evolutionary branching have not yet been investigated with mathematical models. We thus have no theoretical literature to inform predictions against which to compare data. Herein, we propose to investigate the potential effects of complex life-cycles on biological diversification via evolutionary branching by constructing and analyzing a multi-variate evolutionary model of a population of consumers that undergo development in two distinct life-history stages (i.e. juveniles and adults).

We know from the theoretical literature that negative frequency-dependent disruptive selection, which can arise due to the interplay between organisms of a population and their environment, is an important element driving phenotypic diversification. Such selection regime can result from frequency- and density-dependent interactions between the organisms and their environment, so that the fitness landscape itself changes as the population evolves. This can lead to the population reaching a fitness minimum. Fitness minima that are attractors of the evolutionary dynamics are known as evolutionary branching points (Geritz et al., 1998), where a population experiences disruptive selection. Under this regime, more divergent phenotypes are then favored over intermediate ones (Rueffler et al., 2006b), which in turn leads to phenotypic diversification (Dieckmann and Doebeli, 1999; Dieckmann et al., 2004; Bolnick, 2006; Doebeli et al., 2007; Pennings et al., 2008; Ripa, 2009). Branching points occur in many models in which fitness is derived from ecological scenarios that account for resource and mate competition, predation, pathogens, etc.

The phenomenon of evolutionary branching is well understood, under a variety of different ecological scenarios, for simple cases where interactions are mediated by a single quantitative trait in an unstructured life-cycle (Kisdi and Geritz, 1999; Doebeli and Dieckmann, 2000; van Doorn and Weissing, 2002; Rueffler et al., 2006a; Ma and Levin, 2006; Pennings et al., 2008; Zu et al., 2011). More recently, advances have been made in expanding our knowledge under what conditions we can expect evolutionary branching to occur when considering suites of co-evolving traits that interact in their effects on fitness (Doebeli and Ispolatov, 2010; Débarre et al., 2014; Svardal et al., 2014; Geritz et al., 2016; Vasconcelos and Rueffler, 2020). But this too is under the assumption of an organism with a single life-history stage.

However, the developmental trajectories through which organisms move from embryos to adulthood are spectacularly diverse. More than half of the earths biodiversity have a life-cycle which can be classified as complex – that is, one in which individuals undergo some type of ontogenetic niche shift (Hall and Wake, 1999; Wilbur, 1980; Werner and Gilliam, 1984; Werner and Hall, 1988), making this the most widely used life-history strategy among animals (Truman and Riddiford, 1999). For instance, arthropods and fish – two of the groups that include most organisms with this form of complex lifecycle – account for more than 85% of all animal biomass (Bar-On et al., 2018). Myriad organisms – among them insects, fishes, amphibians, mollusks, crustaceans, cnidarians, echinoderms – undergo metamorphosis, the most dramatic of life-cycles. Metamorphosis is an ancestral trait in amphibians, but in insects it is a derived state. It evolved only once from forms that underwent more direct development(Truman and Riddiford, 1999; Truman, 2019), and, in insects, metamorphosis seems to be an irreversible trait, with no known cases of reversions to ametabolous or hemimetabolous development (Yang, 2001). Metamorphosis is radical in that it entails the appearance of different body parts, remodelling of organs and restructuring of body plan between larval and adult forms, who can inhabit environments as disparate as aquatic and terrestrial.

The radically different morphologies in larval and adult stages of metamorphosing animals amount to specific feeding, locomotive, physiological, and behavioral adaptations, which means they occupy distinct ecological niches and interact with different sets of prey and predators. This implies that each life-stage is under discrepant selective pressures, and the empirical evidence points to a sharp decoupling of traits and their evolutionary trajectories across the life-cycle (Wollenberg Valero et al., 2017; Sherratt et al., 2017).

But complex life-cycles can encompass more than the extreme case of metamorphosis. Hemimetabolous insects and some fish (McMenamin and Parichy, 2013), while retaining their body plans throughout life-stages, undergo morphological changes to their feeding apparatuses related to dietary intake. For instance, the nymphs of dragonflies possess a prognathous head with prominent labial hooks adapted for catching aquatic prey. But the adults are flying predators who catch their prey mid-air with their front legs, which are located below their heads. This requires a different adaptation, and their heads develop into a hypognathous orientation so they can bring food up into their mouths (Popham and Bevans, 1979).

Such ubiquity begs for an explanation, which, in evolutionary biology tends to be adaptive. It is widely thought that complex life-cycles, by facilitating trait decoupling among life-stages (Moran, 1994; ten Brink et al., 2019), accrues some important potential benefits: complex life-cycles can (i) reduce resource competition between juveniles and adults (Ebenman, 1987; Truman and Riddiford, 1999); (ii) facilitate colonization of seasonally available (Istock, 1967; Slade and Wassersug, 1975) or newly arisen niches (Rainford et al., 2014) by the different life-stages; and (iii) foster a life-cycle “division of labor” between juveniles focused on growth, and adults focused on dispersal and reproduction, allowing each to optimise their function independent of the other life-stage (Bryant, 1969; Ebenman, 1992; Rolff et al., 2019). These benefits are clearly illustrated for the case of metamorphosis.

As seen in the above exposition, the role of ecology features prominently in some hypotheses about the evolution of complex life-cycles. And due to eco-evolutionary feedbacks, we can wonder: what are the effects of complex life-cycles in ecosystems? The more recent theoretical literature has furnished us with knowledge of the highly complex population dynamical behavior of systems in which an ontogenetic niche shift occurs. Schreiber and Rudolf (2008) give us one of the first detailed explorations of this phenomenon, where they show that, when each life-stage of an organisms with a complex life-cycle uses a different ecological niche for resource acquisition, they effectively couple these ecosystems in complicated ways that defy intuition. For example, by increasing juvenile habitat productivity, juveniles have more energy to invest in maturation, which causes a dramatic increase in the adult population. This, in turn, increases intra-stage competition among adults, who now fail to acquire enough resources to invest in reproduction, leading to a dramatic decrease in juveniles. By increasing or decreasing model parameters such as consumer feeding efficiency, mortality, and resource carrying capacities, alternative stable states appear, where the distribution of the consumer population at equilibrium can abruptly shift from being dominated by juveniles to being dominated by adults, or vice-versa. This occurs because of a positive feedback loop such as the one described above (Nakazawa, 2011, 2015).

Given how widespread complex developmental programs are among most animals, their intricate effects on population and community structures, it is crucial to ask how do these sharp phenotypic and ecological differences between the life-history stages affect macroevolutionary patterns such as phenotypic diversification – and, further along the evolutionary timescale, speciation? As we have mentioned, the interactions between organisms that undergo metamorphosis and their environment are mediated by separate sets of traits in the different life-stages (Wells, 2007), so it is easy to see that ecological opportunity can present itself either to the larval stage, the adult stage, or both simultaneously. Therefore, disruptive selection can, in principle, emerge at each of these life-stages independently.

This begs the question: is the potential for evolutionary branching in such organisms more likely to be driven by interactions at either one of the life-stages, or at both equally? Evolutionary branching could potentially occur in traits that are manifested in strictly distinct stages of the life-history cycle, whereby the following evolutionary outcomes are logically possible: (i) diversification occurring in only one of the life-stages such that the population becomes polymorphic at either the juvenile or the adult stage while all lineages share the same trait at the other life-stage (e.g., different species use different resources during their larval stage but not during their adult stage), (ii) diversification into two lineages occurring simultaneously at both life-stages, such that individuals of one lineage use different resources at both life-stages compared to individuals of the other lineage, (iii) diversification occurring in both life-stages such that no two lineages share the same resources at both life-stages.

So, we ask, what are the potential evolutionary outcomes of disruptive selection in organisms with complex life-cycles and how are they affected by these intricate interactions across trophic levels? We look at the case that disruptive selection is driven by competition where each life-stage competes for two stage-specific resources, under the assumption that there is a trade-off in performance for feeding on each resource. We investigate these questions under the framework of Adaptive Dynamics (Geritz et al., 1998), in which the fitness function of phenotypes is derived from explicit ecological scenarios, taking into account frequency dependent interactions. The juveniles and adults in this model can be thought of as different modules in the sense of holometabolous insects who do not share resources at all, and thus have no inherent genetic or developmental constraints in how they can evolve independently. We hypothesize that complex life-cycles have the potential to lead to evolutionary branching occurring in juveniles and adults independently, which translates to higher diversification rates.

## Model

We first describe a population dynamical model of a consumer species with two lifestages that feeds on two different resources at each of these stages. We then continue by adding mutant consumers to this community and describe how – based on the invasion success of such mutants – the long-term evolutionary dynamics of juvenile and adult resource feeding traits can be inferred.

### Population dynamics

To investigate the set of questions posed above, we consider an extension of the model by Schreiber and Rudolf (2008) in which consumer species undergo an ontogenetic niche shift, meaning the juvenile and adult stages each live in different habitats where they acquire non-overlapping resources. The juveniles, denoted by *J*, feed on two juvenile specific resources, *R*_1,*J*_ and *R*_2,*J*_; and the adults, denoted by *A*, feed on two adult specific resources, *R*_1,*A*_ and *R*_2,*A*_. In the absence of consumers, the resource species on which juvenile and adult consumer feed grow logistically with intrinsic growth rates *r_k,J_* and *r_k,A_* to carrying capacities *K_k,J_* and *K_k,A_* (*k* ∈ {1,2}). Both juvenile and adult consumers deplete resources according to a linear functional response with resource and life-stage specific feeding efficiencies *a_k,J_* and *a_k,A_*. Juvenile individuals mature at a rate proportional to the amount of resources consumed by juveniles where the constant of proportionality is given by the resource specific juvenile conversion efficiencies *c_k,J_*. Similarly, adults give birth at a rate proportional to the amount of resources consumed by adult individuals where the constant of proportionality is given by the resource specific adult conversion efficiencies *c_k,A_*. Finally, adult and juvenile consumer species die at death rate *d_J_* and *d_A_*, respectively. The dynamics of this system is given by

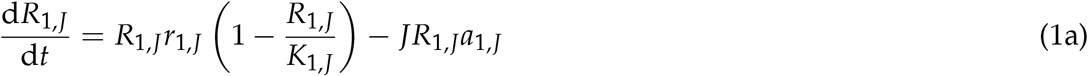

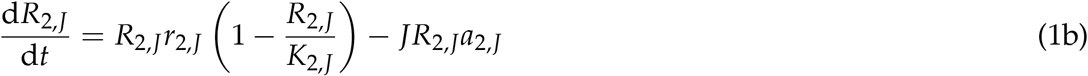

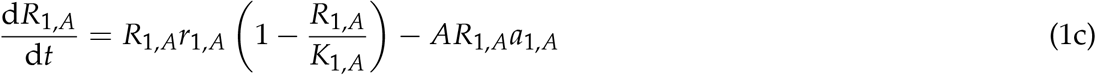

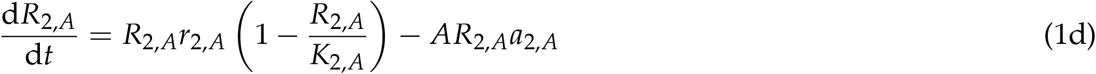

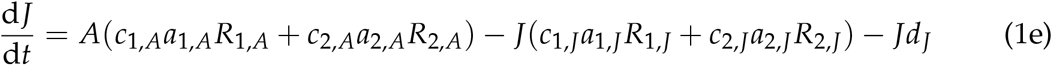

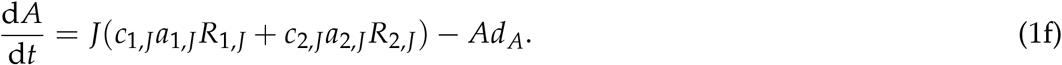

To simplify the population dynamical model we follow previous authors (MacArthur, 1984; Ackermann and Doebeli, 2004) and assume that the dynamics of the resources occur at a faster time-scale than that of the consumers, such that resource densities are always at a quasi-equilibrium determined by the abundance of consumers. This allows us to re-write the dynamics of the consumer species as

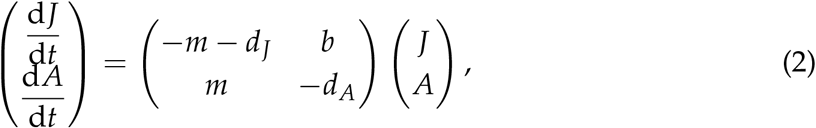

where

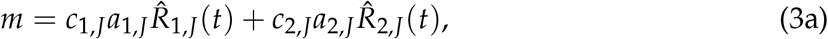

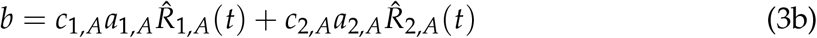

are the maturation and birth rate, respectively. Here, the 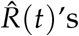 denote the value of the resource quasi-equilibria at time *t*, which are functions of the densities and feeding efficiencies of all consumer species. See Appendix S1 for a full derivation.

Schreiber and Rudolf (2008) showed that this model can exhibit complex population dynamics including limit cycles and alternative stable states. We here restrict ourselves to parameter combinations that result in unique, strictly positive, and stable fixed point equilibria. Depending on model parameters, the population at equilibrium can be dominated by juvenile individuals, adult individuals or the number in the two stages can be equal. We analyze each of these scenarios separately as they qualitatively impinge on our results.

### Evolutionary dynamics

We assume that juveniles and adults can evolve in their feeding efficiencies *a_i,J_* and *a_i,A_*, and that there are stage-specific trade-offs such that performing better at one resource in one life-stage comes at the expense of performing worse at the other resource during the same life-stage. That is, there is a trade-off between the ability to feed on juvenile resource 1 and 2, and another trade-off between the ability to feed on adult resource 1 and 2, but no trade-off between an ability to feed as a juvenile and as an adult. Many organisms with complex life cycles undergo significant changes in their bauplan, and we assume that juvenile and adult traits are coded by independent sets of genes with no pleiotropic effect on the foraging trait at the other life-stage.

The boundary of the set of feasible phenotypes can be described by a trade-off curve. We assume that evolution has reached this constraint and from now on moves along this boundary, which we parametrize as a trade-off curve as shown in Figure 1. Its curvature determines the strength of such trade-off, meaning whether for each unit of gain in performance for one resource there is a small or large loss in performance for the other. This curvature is determined by the parameters *z_J_* and *z_A_* for the juvenile and adult trade-off, respectively. The linear trade-off separates the set of concave tradeoff curves, determined by *z_J_* > 0 and *z_A_* > 0, respectively (weak trade-offs) from the set of convex trade-off curves, determined by *z_J_* < 0 and *z_A_* < 0, respectively (strong trade-offs). We parametrize the trade-off curves in a specialization coefficient, denoted *θ_J_* for juveniles and *θ_A_* for adults, and collect them in the two dimensional trait vector ***θ*** = (*θ_J_*, *θ_A_*). The parametrization of the trade-off curves is such that *θ_J_* = 0 corresponds to a specialist for juvenile resource 1, *θ_J_* = 1 to a specialist for juvenile resource 2, and *θ_J_* = 0.5 to the generalist that is equally specialized for both juvenile resources, and the same applies to the adult trait *θ_A_* and adult resources. We consider these two specialization coefficients as the evolving traits in each life-stage, so that *a*_1,*J*_ and *a*_2,*J*_ are functions of *θ_J_*, and *a*_1,*A*_ and *a*_2,*A*_ are functions of *θ_A_*. In contrast to many other studies using trade-off curves (e.g. Levins, 1962; Rueffler et al., 2006a; Vasconcelos and Rueffler, 2020) we parametrize the trade-off curve such that feeding efficiency for the generalist is independent of the trade-off curvature (all curves in Figure 1 pass through the same point). The reason is that for our purpose it is useful to be able to vary the curvature of the trade-off curve without affecting the values of the population dynamic equilibria of a generalist consumer species. For the formula used to parametrize the trade-offs we refer to Appendix S2.

**Figure 1:**
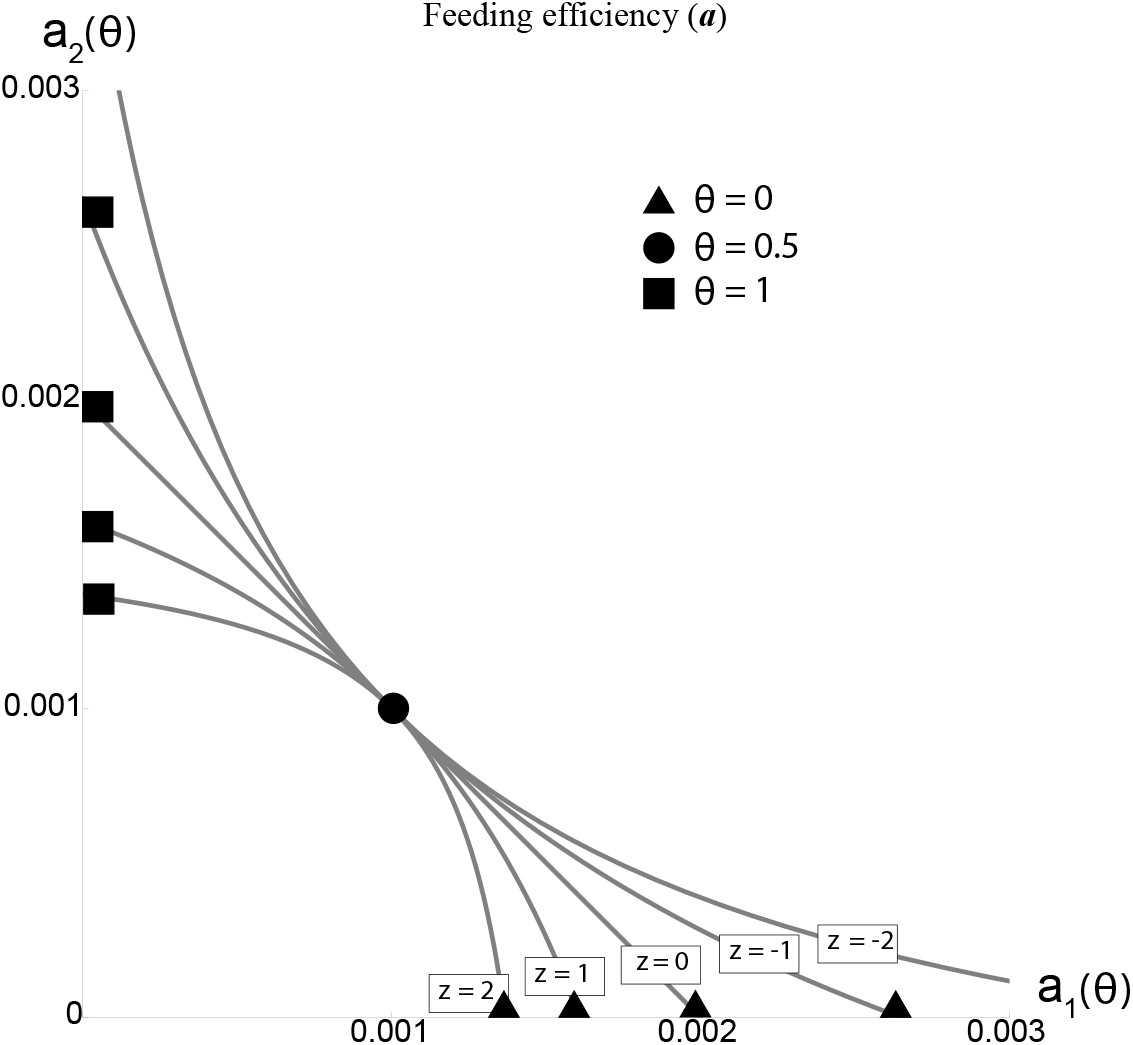
Trade-off curves for the two resource specific feeding efficiencies as determined by equation (S8). The same trade-off parametrization is used for both the two juvenile and the two adult feeding efficiencies and the subscripts *J* or *A* are omitted here for brevity. Trade-off curves are parametrized such that *θ* = 0 corresponds to a specialist for resource 1 (triangle), *θ* = 1 to a specialist for resource 2 (square), and *θ* = 0.5 to the generalist that is equally specialized for both resources 1 and 2 (circle). The parameter *z* gives the strength of the trade-off, where *z* < 0 corresponds to a strong trade-off and *z* > 0 corresponds to a weak trade-off. The generalist strategy is fixed at *a*_gen_ = 0.001 regardless of trade-off curvature, which allows us to investigate the effects of varying the trade-off geometry without concern for unwanted effects on population composition and dynamical stability.

We study the evolutionary trajectories of the traits *θ_J_* and *θ_A_* using the framework of adaptive dynamics (Metz et al., 1996; Geritz et al., 1998), which is based on the technical assumptions rare mutations occurring in very large population. The former assumption implies that resident populations reach their demographic attractor before a new mutant appears, while the latter assumption allows us to ignore the possibility that deleterious mutations increase in frequency due to drift. For simplicity, we assume clonal organisms, an assumption that does not affect the course of the monomorphic evolutionary dynamics. Individual based simulations have shown that the conclusions derived from this framework are usually robust to violations of its assumptions (Champagnat et al., 2006; van Doorn et al., 2009; Svardal et al., 2014; Vasconcelos and Rueffler, 2020). We denote the resident’s trait values 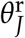 and 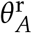, and the mutant’s 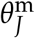 and 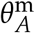.

The population dynamics of a mutant sub-population when rare is given by

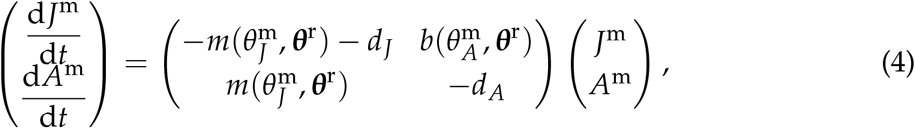

where

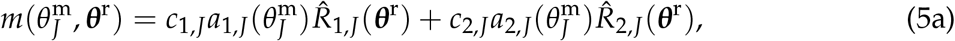

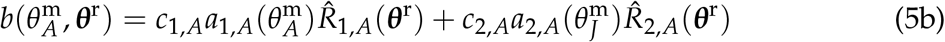

are the maturation and birth rate, respectively. Here, 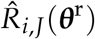 and 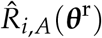 denote the densities of the resources at equilibrium of equation (1) for a resident species with trait vector 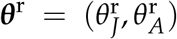. Note that the functions *m* and *b* depend on the traits of both mutant and resident, because the resource densities at equilibrium are determined by the resident phenotype, while the mutant’s phenotype determines its ability to perform in the environment set by such resident.

The dominant eigenvalue of the matrix on the right-hand side of equation (4), denoted by *w*(***θ***^m^, ***θ***^r^), describes the mutant’s expected *per capita* growth rate while rare in a resident population at equilibrium. Its expression is given by equation (S13) in Appendix S3. This growth rate is called invasion fitness (Metz et al., 1992). If *w*(***θ***^m^, ***θ***^r^) < 0, then the mutant will certainly go extinct. More interestingly, the mutant has a positive probability of invading the resident population if *w*(***θ***^m^, ***θ***^r^) > 0. Furthermore, if the mutant’s trait vector ***θ***^m^ is sufficiently close to the resident’s trait vector ***θ***^r^, then successful invasion implies that the mutant will ultimately replace the resident (Dercole and Rinaldi, 2008), resulting in a trait substitution.

Adaptive dynamics (Metz et al., 1996; Dieckmann and Law, 1996; Geritz et al., 1998) is a method to study the dynamics of such trait substitution sequences. The direction of evolutionary change depends both on the mutations occurring in a population and on the direction in trait space showing the steepest increase of the fitness landscape, given by the selection gradient 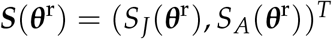 with components

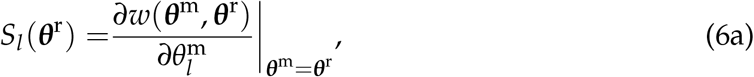

where *l* stands for *J* and *A*. Trait vectors 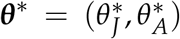 that make both components of ***S***(***θ***^r^) zero simultaneously are referred to as singular points and we denote them with an asterisk superscript. In multidimensional trait spaces a full classification of the possible evolutionary dynamics in the neighborhood of singular points is in general complex due to the dual dependence on the mutational input and the fitness landscape (Leimar, 2009; Geritz et al., 2016; Vasconcelos and Rueffler, 2020). However, for our model this classification is in fact straightforward since, as our analysis shows, it is not affected by the details of the mutational input (see Appendix S3 for details). Biologically speaking, the reason is that the effect on fitness of mutations changing the juvenile trait are independent of mutations changing the adult trait. At singular points these traits show no pleiotropic interactions in the sense of Débarre et al. (2014).

In our model, singular points ***θ\**** are of one of three types. First, if ***θ\**** is an attractor of the evolutionary dynamics (also called convergence stable) and no nearby mutant can invade a resident population characterized by the singular strategy, then ***θ***\* is an endpoint of the evolutionary dynamics. Second, if ***θ***\* is convergence stable but invadable by some nearby mutants, then ***θ***\* is called an evolutionary branching point. This is the case we are most interested in. At such points, mutants do not replace the residents with the singular trait value, but can coexist with them, and the population changes from monomorphic to dimorphic. Each lineage then experiences selection in an opposing direction, resulting in increased phenotypic divergence. Third, a singular point can be a repellor of the evolutionary dynamics in which case it is never reached by gradual evolution.

We analyze the evolutionary dynamics of the traits *θ_J_* and *θ_A_* in two steps. First, we investigate the evolutionary dynamics of these traits independent of each other by keeping one of the traits fixed and determine the location and evolutionary properties of the singular points for the other trait. This analysis, which is based on the analytical and numerical calculations described in the Appendices S3 and S4, allows us to compare the conditions for evolutionary diversification in the two life-stages.

In the second analysis, we study the co-evolutionary dynamics of *θ_J_* and *θ_A_*. The focus is to investigate the final phenotypic composition of evolved communities. This analysis is based on stochastic computer simulations (whose algorithm is described in Appendix S5) that take into account that the direction and speed of evolution of different lineages is affected by demographic stochasticity during the establishment of new mutants.

For simplicity, we focus on symmetric parameter values such that 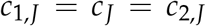, 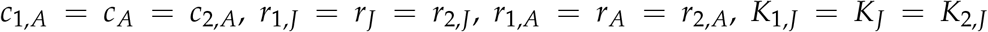, and 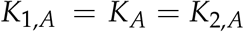. We do, however, also investigate a case in which allow for asymmetry in the resource growth rates.

Full symmetry enforces that the trait vector 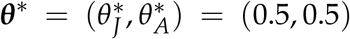, for which the consumer population is a perfect generalist for both resources at both life-stages, is a singular point. We investigate the evolutionary dynamics at this singular point for three different sets of parameters, resulting in a consumer population (i) dominated by juvenile individuals, (ii) dominated by adult individuals, and (ii) consisting of an equal number of juvenile and adult individuals. We use the parameters in Table 1 to obtain the different population dynamical regimes, resulting in a ratio of juvenile to adult individuals at equilibrium at the generalist singular point equal to 2.4 and 0.43 for the juvenile and adult dominated case, respectively.

**Table 1:** Parameter values used to produce Figures 2–5.

|  | Juvenile | Adult | Equal |
| --- | --- | --- | --- |
| $c_J$ | 0.25 | 0.5 | 0.25 |
| $c_A$ | 0.5 | 0.5 | 0.5 |
| $d_J$ | 0.6 | 1 | 1 |
| $d_A$ | 1 | 1 | 1 |
| $K_J$ | 1500 | 3000 | 3000 |
| $K_A$ | 3000 | 3000 | 3000 |
| $r_J$ | 6 | 6 | 6 |
| $r_{1,J}$ | 4 | 4 | 4 |
| $r_{2,J}$ | 8 | 8 | 8 |
| $r_A$ | 6 | 6 | 6 |
| $r_{1,A}$ | 4 | 4 | 4 |
| $r_{2,A}$ | 8 | 8 | 8 |

## Results

### One-dimensional evolutionary dynamics – The symmetric case

In this section, we analyze the evolutionary dynamics of each trait under the thought experiment that the other trait is fixed at the generalist trait value *θ* = 0.5. Figure 2 shows bifurcation diagrams for the singular points of each trait as a function of the corresponding trade-off curvature. The evolutionary dynamics when restricted to one trait at a time are largely in agreement with the results of comparable earlier studies investigating single trait evolution (Ma and Levin, 2006; Rueffler et al., 2006a; Zu et al., 2011; Vasconcelos and Rueffler, 2020).

**Figure 2:**
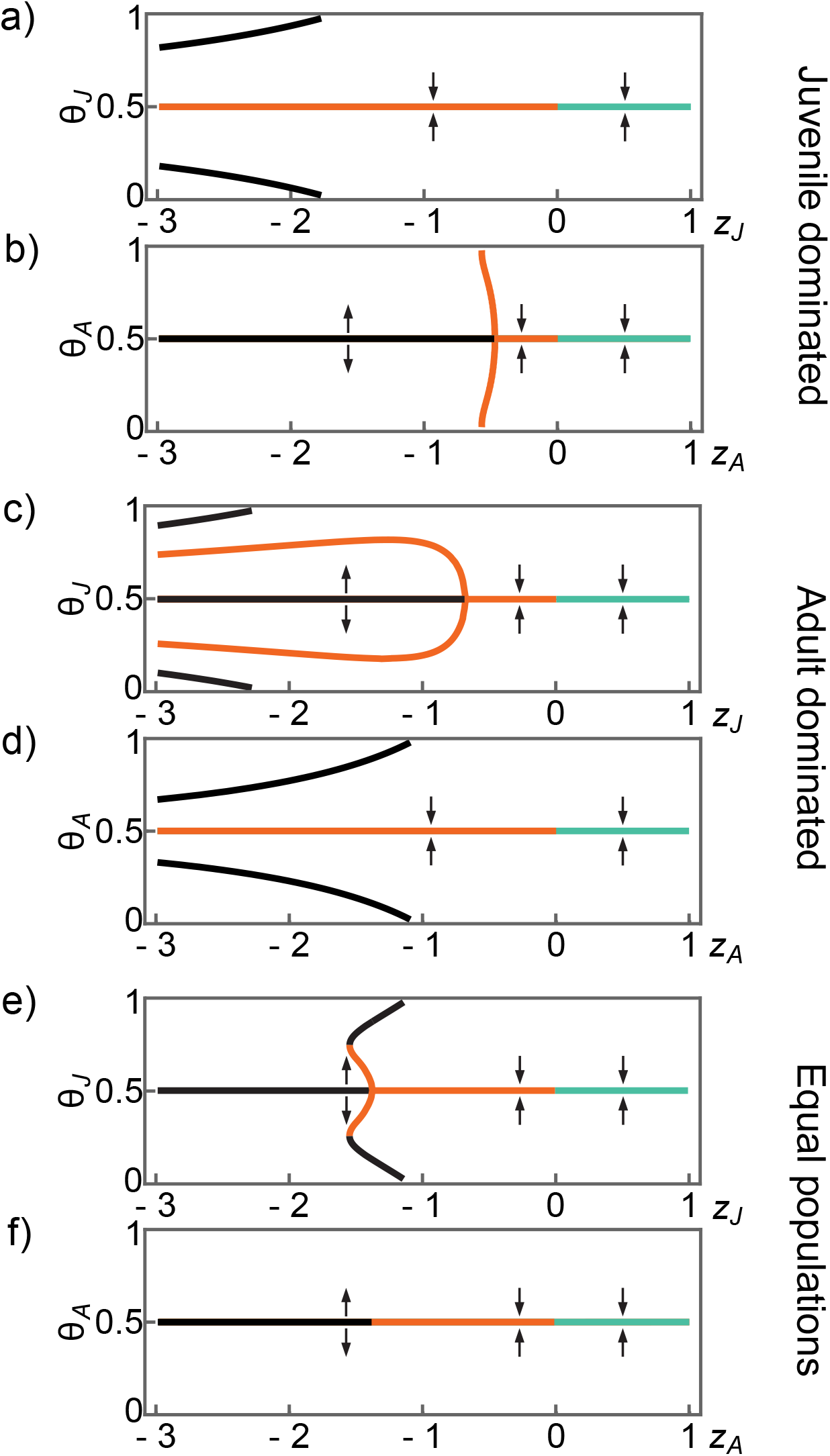
Bifurcation diagrams for the case that only the juvenile or only the adult trait evolves while the non-evolving trait is fixed at the generalist strategy. Parameters (see Table 1) are symmetric within each life-stage and give rise to a juvenile dominated population in panel (a) and (b), to an adult dominated population in panel (c) and (d) and to equal population sizes for juveniles and adults in panel (e) and (f). The x-axis gives the curvature parameter *z_J_* in panels (a), (c) and (e), and the curvature parameter *z_A_* in panels (b), (d), and (f). The y-axis gives the trait space of the evolving trait *θ_J_* in panels (a), (c) and (e), and the trait space of the evolving trait *θ_A_* in panels (b), (d) and (f). The location of singular points is shown by colored lines, where black corresponds to a repelling singular point, orange corresponds to an evolutionary branching point, and green corresponds to an uninvadable attracting singular point. Arrows indicate the direction of the selection gradient.

First, due to symmetry the generalist trait value is always a singular point, but more than one singular point can exist for strong trade-offs (*z* < 0). Second, the generalist singular point is always an evolutionary endpoint for weak trade-offs (*z* > 0). Third, singular points for strong trade-offs are always invadable. Fourth, the generalist singular point changes from an evolutionary endpoint to an evolutionary branching point where trade-offs change from weak to strong (at *z* = 0). When evolutionary branching occurs the two emerging lineages evolve to become perfect specialists for the two different resources. Fifth, the generalist singular point can change from an evolutionary branching point to an evolutionary repellor for very strong trade-offs (*z* << 0).

Importantly, when comparing the bifurcation diagrams for our different demographic scenarios, we find that the interval for which the generalist singular point is an evolutionary branching point is larger for the trait corresponding to the more abundant life-stage (compare figure 2a&b with c&d).

Indeed, in Appendix S4 we prove that the interval of trade-off curvatures for which the generalist singular point is an evolutionary branching point is larger for the adult trait than for the juvenile trait if the ratio of adult to juvenile equilibrium population size is larger than the ratio of adult to juvenile resource growth rate, 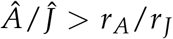, and smaller if the opposite inequality holds true (note that we assume *r_J_* = *r_A_* in Table 1). As a corollary, when both juvenile and adult population densities are identical, evolutionary branching occurs for identical intervals of trade-off curvatures in the two traits (figure 2e&f). The fact that the trait of the more abundant life-stage always has a larger interval of trade-off curvatures that lead to evolutionary branching shows that stronger competition favors diversification.

We also find patterns that were not known from previous theoretical studies of resource specialization for two discrete resources. For some intervals of convex trade-off curvatures up to five singular points exist simultaneously (figure 2c&e), while for other intervals a repelling generalist singular point is flanked on each side by a branching point (figure 2b,c&e). In the latter case, populations evolve away from the generalist strategy and become dimorphic at a singular point that corresponds to incomplete specialization for one of the resources.

### Two-dimensional evolutionary dynamics – The symmetric case

Figure 3 shows the simulated evolutionary dynamics for the co-evolving juvenile and adult trait values under different population dynamical regimes for different trade-off curvatures and symmetric parameter values as shown in Table 1. The focus of this analysis is to study the pattern of evolved communities after evolutionary branching. Under symmetric parameter values, we find that at most one branching event takes place resulting in two coexisting phenotypes. It is easy to show that on an ecological time scale three phenotypes can coexist, but results by Saltini et al. (2022) show that such communities are not attainable through gradual evolution and other processes such as immigration have to be invoked.

**Figure 3:**
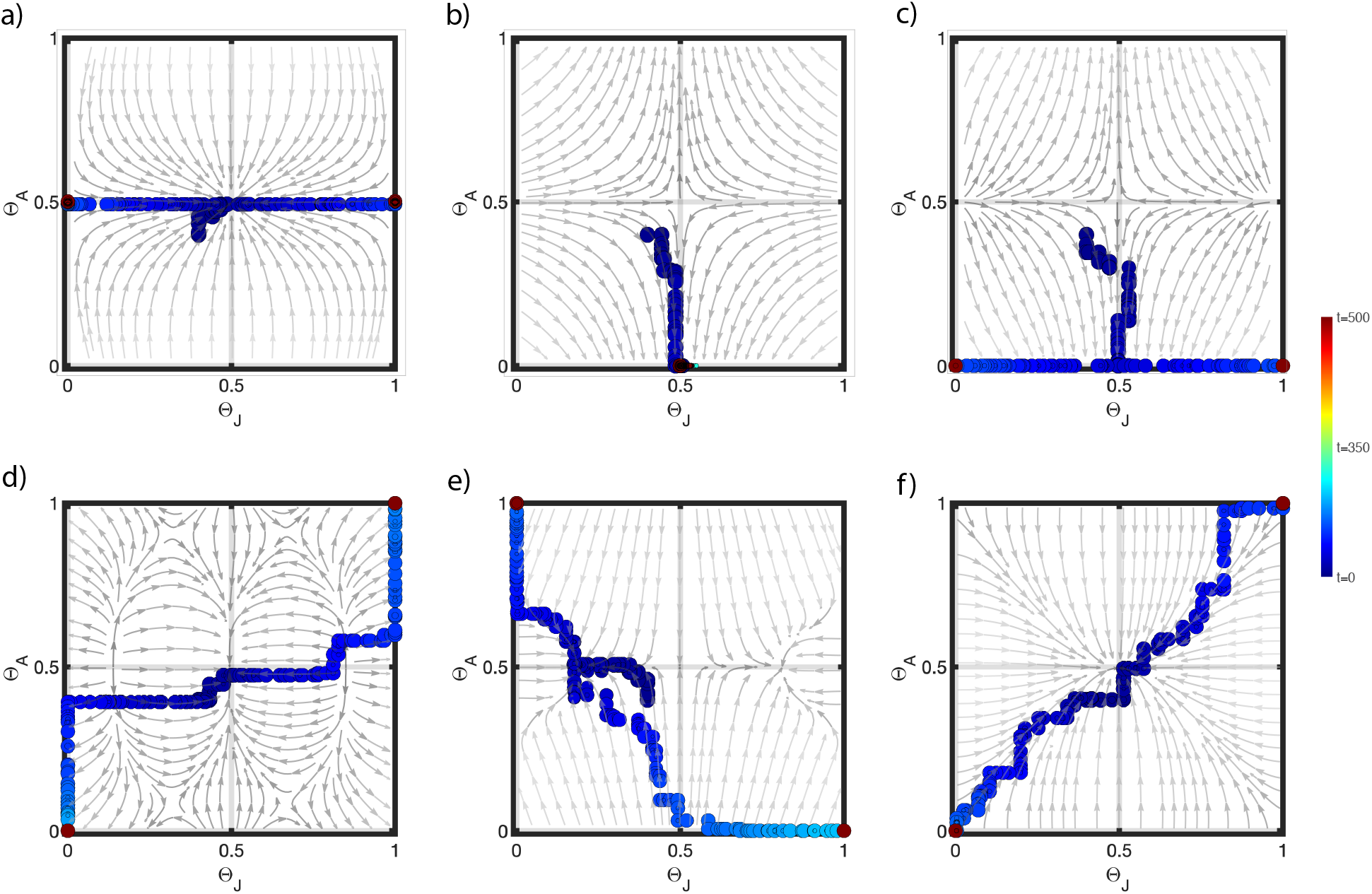
Evolution in the (*θ_J_*, *θ_A_*) phase-plane under different population dynamical regimes and for different trade-off curvatures. All parameters are symmetric as specified in Table 1. a) Juvenile dominated population with *z_J_* = −1 and *z_A_* = 0.5. b) Juvenile dominated population with *z_J_* = 0.5 and *z_A_* = −1.5. c) Juvenile dominated population with *z_J_* = −1 and *z_A_* = −1.5. (d) Juvenile dominated population with *z_J_* = −1.5 and *z_A_* = −0.4. (e) Adult dominated population with *z_J_* = −1.5 and *z_A_* = −0.5. (f) Population consisting of an equal number of juveniles and adults with *z_J_* = −0.5 and *z_A_* = −0.5. Gray arrows indicated the direction of selection for a monomorphic population as given by equation (S15) and colored lines show simulated evolutionary trajectories starting from a population with trait vector (*θ_J_*, *θ_A_*) = (0.4,0.4). Each circle represents a population in time after a successful mutation event, and colors indicate the passage of time counted as successful mutations where dark blue is *t* = 0 and dark red is *t* = 500.

For certain combinations of trade-off curvatures we can infer the outcome of the co-evolutionary dynamics of the juvenile and adult trait from the analysis of the independently evolving trait presented in the previous section. First, if both trade-offs are weak 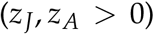, then 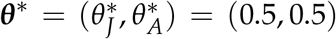 is a unique singular point. This singular point is both convergence stable and uninvadable, and therefore the population is expected to eventually consist of a single species that is a perfect generalist at both the juvenile and adult stages. This outcome is evident from the fact that evolution of each trait in isolation is predicted to evolve to the generalist trait value (green lines in Figure 2) and that the two traits do not interact in their effect on fitness (as shown in Appendix S3). This is in contrast to results by Vasconcelos and Rueffler (2020, Chapter 1) who report that a co-evolutionary singular point can be a branching point even if both traits in isolation evolve to an uninvadable singular point (compare their Figure 4 and 5e).

**Figure 4:**
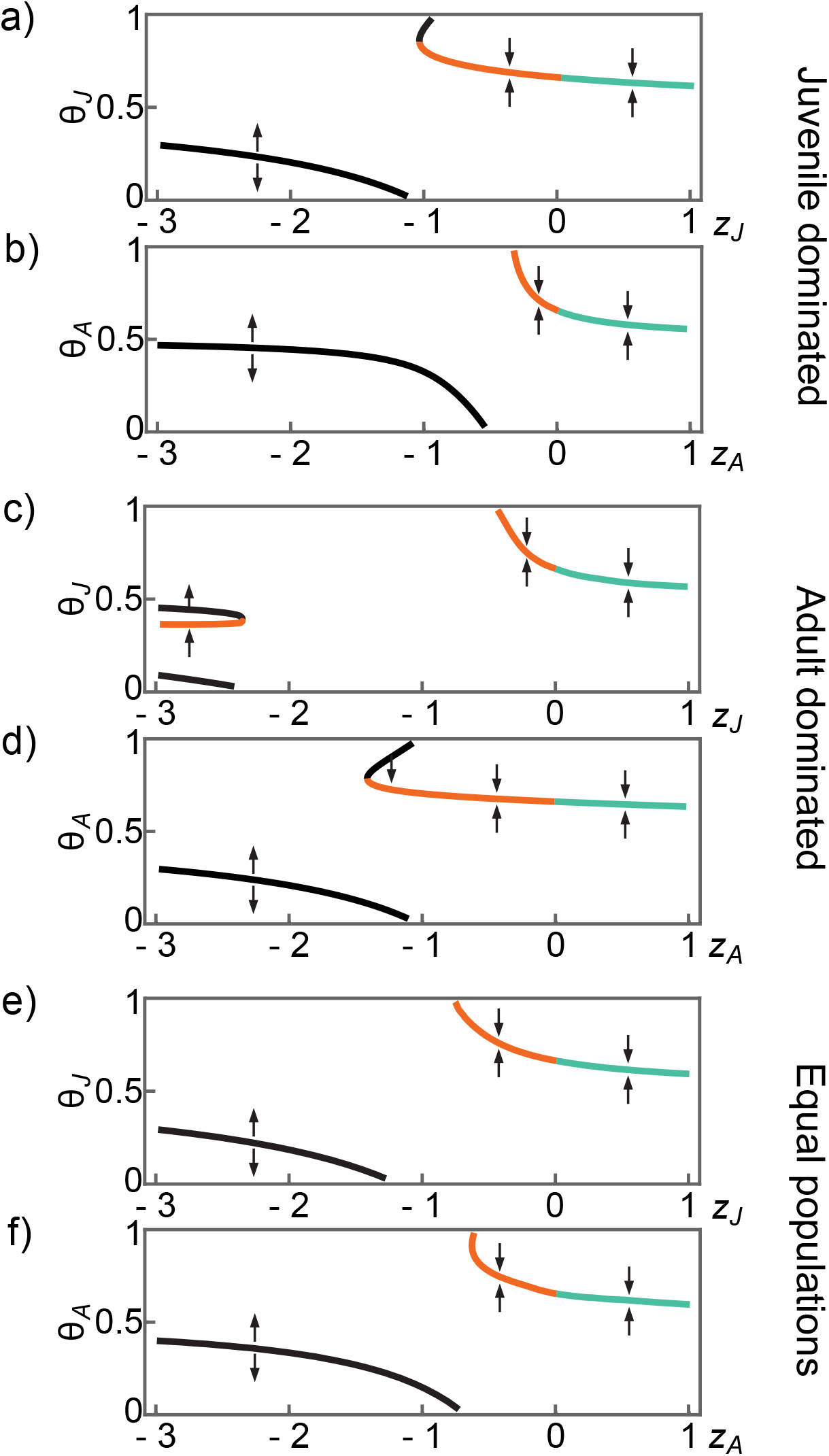
Bifurcation diagrams analogous to Figure 2 but now with asymmetry in the resource growth rates (*r*_1,*J*_ < *r*_2,*J*_ and *r*_1,*A*_ < *r*_2,*A*_).

**Figure 5:**
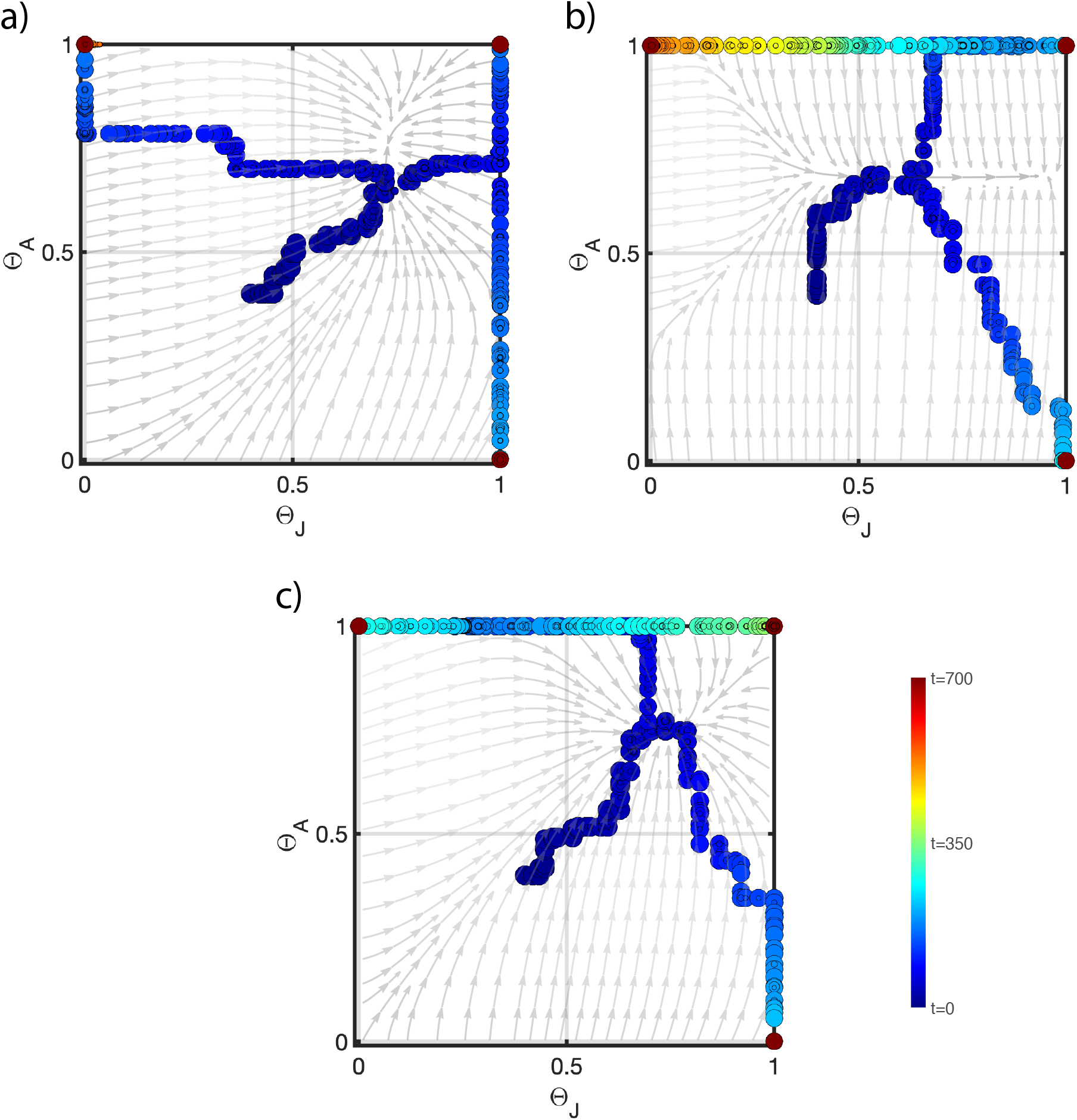
Evolution in the (*θ_J_*, *θ_A_*) phase-plane under different population dynamical regimes and for different trade-off curvatures. Within stage resource growth rates are asymmetric as specified in Table 1. (a) Juvenile dominated population with *z_J_* = −0.8 and *z_A_* = −0.2. (b) Adult dominated population with *z_J_* = −0.5 and *z_A_* = −0.5. (c) Population consisting of an equal number of juveniles and adults with *z_J_* = −0.5 and *z_A_* = −0.5. Gray arrows indicated the direction of selection for a monomorphic population as given by equation (S15) and colored lines show simulated evolutionary trajectories starting from a population with trait vector (*θ_J_*, *θ_A_*) = (0.4,0.4). Each circle represents a population in time after a successful mutation event, and colors indicate the passage of time counted as successful mutations where dark blue is *t* = 0 and dark red is *t* = 700.

Second, when a weak trade-off in one trait is combined with a strong trade-off in the other trait, such that the trait with the strong trade-off approaches a branching point when evolving in isolation (for instance, Figure 2a&b for *z_J_* < 0 and *z_A_* > 0), then evolutionary branching occurs in one life-stage while individuals in the other life-stage all adopt the generalist trait value. Thus, the population becomes polymorphic at only one of the two life-stages (Fig 3a). This outcome is evident because convergence stability and uninvadability at the generalist of the trait corresponding to the weak trade-off is unaffected by the evolutionary dynamics at the other trait (equations S18 and S20 in Appendix S3).

Third, when a weak trade-off in one trait is combined with a strong trade-off in the other trait such that the trait with the strong trade-off has a single repelling singular point (for instance, Figure 2a&b for *z_J_* > 0 and *z_A_* < 0.6), then evolution results in a single species that is a specialist for one resource at one life-stage and a generalist at the other life-stage (Figure 3b). The reasoning for this outcome is identical to the one given for the previous case.

Fourth, when two strong trade-offs are combined such that one trait evolves toward a branching point while the other trait has only a single repelling singular point (for instance, Figure 2a&b for *z_J_* < 0 and *z_A_* < 0.6), then the outcome can depend on the specifics of the case. In principle, the evolutionary dynamics of one trait can affect whether a singular point for the other trait is an evolutionary branching point or repellor. This can happen since the trait value at one stage affects the other, and evolutionary branching requires strong high population densities resulting in strong competition. Figure 3(c) shows a case where the evolutionary dynamics results in two species, both of which are a specialized for the same resource at one life-stage and specialized for alternative resources at the other life-stage.

The single case that we have not analyzed so far is that of two strong trade-offs such that both traits in isolation evolve toward a branching point. Under this scenario, we always find that a single branching event takes place resulting in two resource specialists for mutually exclusive sets of resources, and therefore always occupy either the diagonal or anti-diagonal corners of the trait plane. In the following we highlight three such cases. Figure 3(d) shows the case for a juvenile dominated population with *z_J_* = −1.5 and *z_A_* = −0.4. Thus, when evolving in isolation both traits have a branching point at the generalist strategy (horizontal orange lines in Figure 2a&b). We observe that the population first evolves to the vicinity of the co-evolutionary singular point ***θ\**** = (0.5,0.5) and then branches in a diagonal direction. Inspecting many simulation runs, we find that branching results equally often in a diagonal and an anti-diagonal trait distribution, determined by the stochasticity inherent in our simulation algorithm at the stage of the establishment of the next successful mutant.

Figure 3(e) shows the case for an adult dominated population with *z_J_* = −1.5 and *z_A_* = −0.5. For these trade-off curvatures, when evolving in isolation, the juvenile trait evolves to one of the branching points that flank the repelling generalist trait value while the adult trait value evolves toward the branching point at the generalist trait value (Figure 2c&d). Since our starting trait vector equals (*θ_J_*, *θ_A_*) = (0.4,0.4), the juvenile trait value initially decreases while the adult trait value initially increases. Once the population has evolved a trait vector approximately equal to (*θ_J_*, *θ_A_*) = (0.7,0.5), diversification occurs first in the direction of the adult trait and later also in the juvenile trait value.

Finally, Figure 3(e) shows the case for a population consisting of an equal number of juveniles and adults. The trade-off curvatures are *z_J_* = −0.5 = *z_A_*. In this case, both traits in isolation evolve to a branching point at the generalist trait value (horizontal orange lines in Figure 2e&f). Repeating simulations results in an equal proportion of runs with a diagonal and an anti-diagonal trait distribution.

### One-dimensional evolutionary dynamics – The asymmetric case

We introduce asymmetry into our model by assuming differences in the resource growth rates such that *r*_1,*J*_ < *r*_2,*J*_ and *r*_1,*A*_ < *r*_2,*A*_ (see Table 1). This asymmetry breaks the symmetry of the bifurcation diagrams (compare Figure 2 and 4) without qualitatively changing the results. The attracting generalist singular trait value of the symmetric case is shifted towards the faster growing resource, resulting in partial specialization for resource 2. Furthermore, the repelling generalist singular trait value is shifted towards the slower growing resource. Taken together, this results in a larger basin of attraction for the evolution of specialists for the faster growing resource 2.

### Two-dimensional evolutionary dynamics – The asymmetric case

The co-evolutionary dynamics of the juvenile and adult trait under asymmetric parameter values can result in qualitatively different evolutionary outcomes not observed under symmetry. When both trade-offs are strong such that each trait in isolation evolves toward an evolutionary branching point, then two branching events resulting in a final community consisting of three species can occur. These species are complete specialists and thus occupy three of the four corners of the phase plane. Under parameters for which such trimorphic communities evolve, they do so in up to half of all simulation runs.

Figure 5 shows three simulations that all result in three coexisting species. The used parameters correspond to (a) a juvenile dominated population with *z_J_* = −0.8 and *z_A_* = −0.2, (b) an adult dominated population with *z_J_* = −0.5 and *z_A_* = −0.5, and (c) a population consisting of an equal number of juvenile and adults individuals (given a resident population with (*θ_J_*, *θ_A_*) = (0.5,0.5)) with *z_J_* = −0.5 and *z_A_* = −0.5. The grey arrows describing the gradient under monomorphic evolution show that in all three cases a single singular trait vector 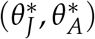 exists, and that this singular point is convergence stable (gray arrows point toward it). The position of these singular points correspond well to those of the analysis for a single evolving trait (that is, the position of the orange line in Figure 4 in the corresponding panel and for the corresponding trade-off curvature). In all three simulations, the evolutionary trajectories start at the trait vector (*θ_J_*, *θ_A_*) = (0.4,0.4) and move in the direction of the singular point. In the juvenile dominated and equal population size case, the monomorphic evolutionary dynamics reaches the vicinity of the singular point where the population branches into two lineages. The situation is different for the adult dominated case (Figure 5b). Here, evolutionary branching occurs in the adult trait once the population has reached the value of the adult singular trait value 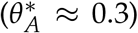 but at a time when the juvenile trait still experiences directional selection toward the juvenile singular trait value 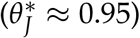. The initial direction of diversification is well predicted by the population composition, that is, in the juvenile trait in the juvenile dominated case, in the adult trait in the adult dominated case, and in a compound direction in the equal population size case.

In all three simulations, the lineage that first reaches the boundary of the trait space undergoes a second branching event and the two lineages resulting from this event undergo character displacement along that boundary. The final outcome is a community consisting of three species. Due to the asymmetry inherent in this model, the final composition of three-species communities is not affected by stochasticity and consistent across simulation runs. This final community composition can be understood as follows. Recall that *r*_1,*J*_ < *r*_2,*J*_ and *r*_1,*A*_ < *r*_2,*A*_. Thus, the species located in the top right corner of the phase plane is specialized for the faster growing resource at both life-stages while the other two species are specialized for the faster growing resource in one life-stage and for the slower growing resource at the other life-stage. The latter two species can coexist with the specialist in the top right corner since each of them has one life-stage in which it uses a resource exploited by no other species. The unoccupied corner corresponds to a hypothetical species being specialized at the slower growing resource at both life-stages. Such a species would not be able to exist in this community since both resources are already used by species that have the advantage of utilizing a faster growing resource at their other life-stage.

## Discussion

In this paper we investigated the effects of complex life-cycles – defined as “a life history that includes an abrupt ontogenetic change in an individual’s morphology, physiology, and behavior, usually associated with a change in habitat” (Wilbur, 1980) – on the potential for biological diversification via gradual evolution and evolutionary branching. We found that life-cycle complexity is able to foster biological diversification via evolutionary branching due to the fact that the life-stage with higher population density is often under a regime of disruptive selection, leading to divergent phenotypic changes related to resource specialization.

Our model can be used as a framework to understand the diversification patterns of organisms with complete metamorphosis, such as amphibians and holometabolous insects, since we assume that the traits of juveniles and adults are determined by independent and non-overlapping genetic mechanisms. This assumption is plausible when referring to organisms that undergo metamorphosis. For example, Johansson et al. (2010) found that while morphology and performance were correlated within larval and adult stages of the frog *Rana temporaria*, performances were completely uncorrelated between larvae and adults. This is supported by later findings of Wollenberg Valero et al. (2017), who observed that genes related to the development of different morphological structures in the frog species *Xenopus laevis* and *Mantidactylus betsileanus* are expressed in a highly phase-specific pattern, pointing to an uncoupling of phenotypic evolution between tadpoles and adults. Even more striking, Sherratt et al. (2017) found that many species of Australian adult frogs and their tadpoles are, as they put it, “evolving independently”, indicating that they are under the influence of different selection regimes. The tadpole and frog morphologies display contrasting evolutionary trajectories, with the first showing highly convergent evolution and low levels of phylogenetic signal, in opposition to the latter, where one can clearly discern clades.

Our initial hypothesis that complex life-cycles lead to higher diversification compared to simple life cycles was partly confirmed. We found that, in the fully symmetric case, there can only be one evolutionary branching event, such that gradual evolution cannot lead to more than two coexisting specialist phenotypes. Saltini et al. (2022) found something similar in their explorations of community assembly. They found that many more species can in principle coexist in trait space, but such communities are not attainable through gradual evolution due to the fact that fitness peaks in the dynamic fitness landscape appear in distant locations. Two morphs competing for, e.g., the same juvenile resource, will produce fewer adults through maturation than if each used a different juvenile resource. The same argument holds true for the adult stage, that is, two morphs competing for the same adult resource will produce fewer progeny than if each used a different adult resource. Thus, evolution favors minimizing intra-stage resource competition, thereby leading to configurations in the trait plane where specialists do not share resources within a life-stage. The final combination of specialists (whether arranged diagonally or anti-diagonally in trait space) is stochastically determined by the sequence of mutation steps. The aforementioned gives evidence that evolution favors character displacement as a way to minimize overlap in resource sharing.

In the asymmetric case, however, due to differences in resource growth rates, we find that gradual evolution can lead to two separate evolutionary branching events, resulting in three coexisting specialist phenotypes. With the introduced asymmetry in within-stage resource growth, a type that feeds on, e.g., a juvenile resource with lower growth rate can make up for that by, as an adult, feeding on a resource with higher growth rate, thus mitigating the effects of intra-stage competition at another stage. Since complete symmetry in all relevant parameters is an artificial situation, we can expect that three coexisting species occur much more often. The unoccupied lower left corner of the trait plane corresponds to the specialist whose juvenile and adult resources have the lowest combination of intrinsic growth. This means that this specialist cannot compete with the two other specialists with whom it would share juvenile resource 1 or adult resource 1, as it cannot make up for feeding on a worse resource at one stage by feeding on a better resource at another stage. This is the reason why evolutionary branching via gradual evolution does not lead to a four coexisting species – although these four specialist phenotypes can coexist ecologically, meaning that a community of four specialists can be assembled by a process of migration (Saltini et al., 2022).

Regardless of whether evolution in organisms with complex life-cycles leads to more than one evolutionary branching event, the fact that the dominant life-stage has much larger potential for branching seems indeed confirmatory that trait modularity favors diversification. Without trait modularity resulting from complex life-cycles, only a small subset of moderately strong trade-offs leads to evolutionary branching (Rueffler et al., 2006a; Vasconcelos and Rueffler, 2020), the remaining trade-off strengths lead to either one specialist or one generalist. But with complex life-cycles, the life-stage that dominates the population will undergo evolutionary branching under any strong trade-off, leading to two specialist phenotypes. And, with asymmetries in resource growth, there can be a second branching event, leading to three phenotypes. Therefore, it is reasonable to infer that the occurrence of double evolutionary branching events is fairly common.

That strong trade-offs in resource use lead to divergent natural selection has been documented in the empirical literature and, importantly, in organisms which undergo ontogenetic niche shifts. For instance, Westneat (1994) documented a trade-off between speed and force in labrid fishes feeding on either evasive fish prey or hard-shelled mollusk prey, which translated to morphological and functional differences of the jaw. Svanbäck and Eklöv (2003) also found trade-offs in feeding performance of perch which relates their body size and shape to the prey type (benthic vs pelagic) that they feed on. And Konuma and Chiba (2007) found strong trade-offs in feeding efficiency of the malacophagous beetle *Damaster blaptoides*, who have two distinct morphotypes – one with an elongated small head specialized in eating snails with thick shells and wide apertures, and another with a stout large head specialized in crushing snails with soft shells and small apertures.

Consistent with our model, there is also empirical evidence pointing to a positive relationship between trait decoupling among life-stages – what some authors refer to as *modularity* – and diversification. Such trait decoupling is accomplished with utmost efficiency in metamorphosis, such that modularity is used as a proxy for life-cycle complexity. Yang (2001) made a forceful case that modularity facilitates evolvability and adaptive radiations. By comparing, according to four conceptually precise criteria, the historical evolutionary patterns between the sister clades Eu- and Holometabola, he found that the Holometabola had significantly higher rates of diversification than the less modular Eumetabola (a group within the paraphyletic Hemimetabola). His hypothesis also predicted that higher modularity entails higher trait variation, for which some supporting evidence is the presence, in the Holometabola, of much more diverse mouthpart types compared to all of the Hemimetabola (Labandeira, 1997). And Rainford et al. (2014), analyzing phylogenies of hexapod families, found that indeed an increase in the rate of diversification was associated with the arising of complete metamorphosis, further strengthening this hypothesis. In addition, they also showed that radiations within this group are linked to the emergence and radiation of the angiosperms. Their findings support the causal roles of both (i) metamorphosis as a key developmental innovation, and (ii) evolutionary responses to newly arisen ecological opportunities as critical ingredients driving adaptive diversification within the Hexapoda. And, despite coming to a different conclusion, Condamine et al. (2016) did a joint analysis of fossil and molecular data where they found a notable increase in diversification rates of insects which clearly postdates the origin of complete metamorphosis, where most of these shifts were found within the holometabolous clades containing the richest orders.

To the best of our knowledge, this is one of the few theoretical explorations of the role of complex life-cycles on biological diversification – but see Saltini et al. (2022, Chapter 4) and ten Brink and Seehausen (2022). The importance of understanding this phenomenon cannot be overstated, as the majority of life on Earth undergoes developmental programs that encompass niche shifts throughout their life-spans. We found that the life-stage that dominates the population dynamical regime experiences disruptive selection whenever trade-offs are strong, which means this life-stage has a much higher chance of undergoing evolutionary branching. We also found that asymmetries in model parameters are able to foster the appearance of another evolutionary branching event, leading to three coexisting phenotypes. Therefore, we can conclude that life-cycle complexity, in general, favors biological diversification.

## Acknowledgments

This work was funded by a doctoral grant to P.V. from CAPES (Coordenação de Aperfeiçoamento de Pessoal de Nível Superior - Brasil).

## Appendix

### S1 Population dynamics

Equation (1) describes the population dynamics of a single consumer species with a structured life cycle consisting of juveniles and adults and where individuals in each stage forage on two different resources. This population dynamical model can be generalized to include *n* consumer species and then reads

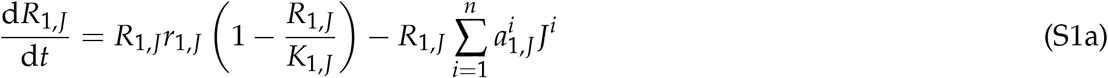

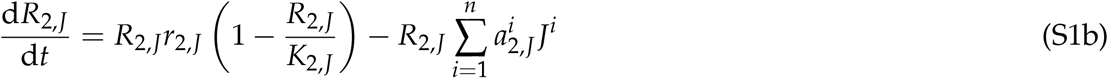

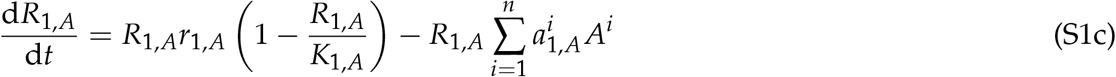

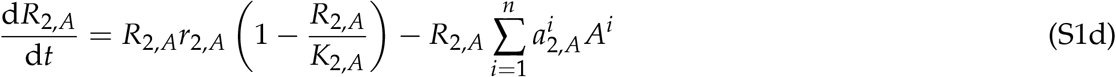

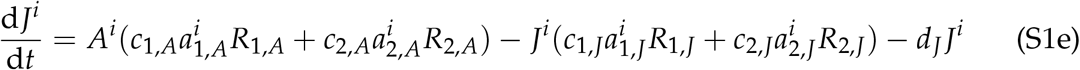

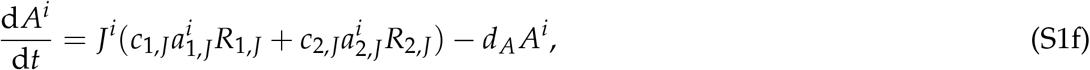

where 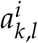 denotes the feeding efficiency of a consumer individual of species *i* ∈ {1, …, *n*}) being is in life-stage *l* ∈ {*J, A*}) for resource species *k* ∈ {1,2} available at that life-stage.

After assuming a time-scale separation between fast resource and slow consumer dynamics the dynamics of the *i*th consumer species can be written in matrix notation as

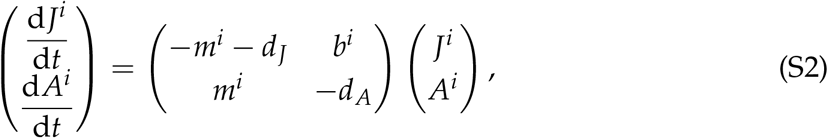

where

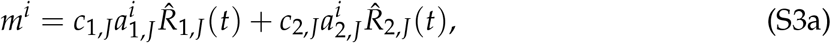

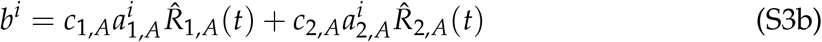

are the maturation and birth rate, respectively, and

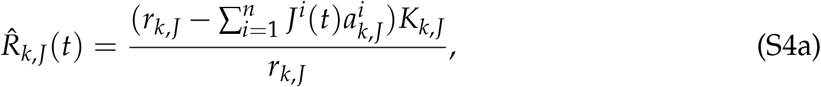

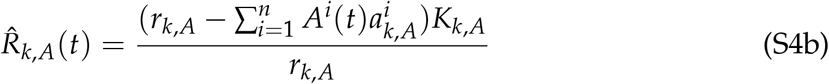

denote the quasi-equilibria for *k*th resource consumed by juveniles and adults, respectively, at time *t*. Note that these quasi equilibria are functions of the densities and feeding efficiencies of all consumer species.

Solving equation (S2) to obtain the consumer equilibrium densities 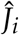 and 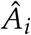 is possible only for monomorphic consumer populations (*n* = 1). That system has, depending on parameters, either one or three positive solutions (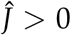 and 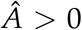), the expressions of which are prohibitively long and we do not present them here. Schreiber and Rudolf (2008) present an analysis of this monomorphic model, showing that it can exhibit a rich dynamics including limit cycles and bistability. For our analysis, however, we restrict ourselves to parameters resulting in a single positive, asymptotically stable point equilibrium. Whenever we need to compute the equilibrium of equation (S2) for more than one consumer species we resort to numerical integration.

### S2 Traits and the trade-off curve

In this study, we analyze the evolutionary dynamic of a trait affecting the two feeding efficiencies of juvenile individuals and another trait affecting the two feeding efficiencies of adult individuals. These traits are assumed to be quantitative morphological or physiological traits that can vary continuously. Each species *i* can then be characterized by a two-dimensional trait vector 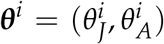, whose entries are the trait values of juvenile and adult individuals, respectively. Here, we describe the functions 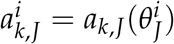 and 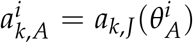 that map stage-specific trait values to the two stage-specific feeding efficiencies.

Our mapping is based on the assumption that the feeding efficiencies at each lifestage are coupled by a trade-off such that a change in the value of a trait resulting in an increase in feeding efficiency for one resource at the corresponding life-stage comes at the expense of performing worse at the other resource at the same life-stage. This can be implemented by assuming that evolution moves along a trade-off curve. As a parametrization of such a trade-off curve we use a formula by Nurmi and Parvinen (2013), which we adapt for our purpose. In the following we omit the subscript indicating species identity and, since we use the identical trade-off parametrization for both the juvenile and adult traits, also skip the subscript for the life-stage.

The parametrization by Nurmi and Parvinen (2013) is given by

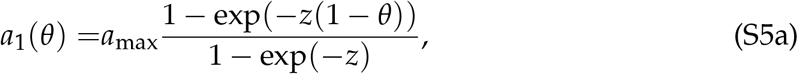

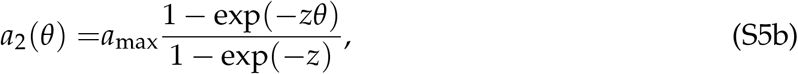

where *a*_max_ is the maximum feeding efficiency of a resource specialist, here assumed to be identical for resource 1 and 2. With this parametrization, the points where the trade-off curve connects to the x- and y-axis are constant while the feeding efficiencies *a*_1_(*θ* = 0.5) and *a*_2_(*θ* = 0.5) of a generalist consumer change as the curvature of the trade-off curve is varied (see figure 1 in Vasconcelos and Rueffler, 2020).

For our purpose, it is advantageous to be able to vary the curvature of the trade-off without changing the values of the feeding efficiencies of a generalist consumer population (see figure 1) and thereby keep the values of its population dynamical equilibrium constant. This can be achieved by making the parameter *a*_max_ a function of the trade-off curvature *z*. To find this function, we use *a*_1_ (0.5) = *a*_gen_ = *a*_2_ (0.5) and solve

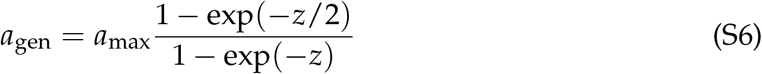

for *a*_max_, resulting in

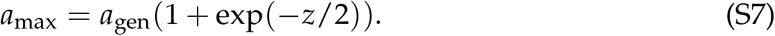

Substituting *a*_max_ in equation (S5) with the right-hand side of equation (S7) gives

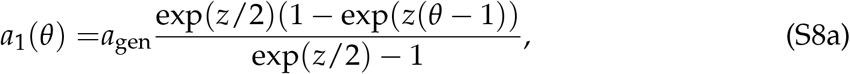

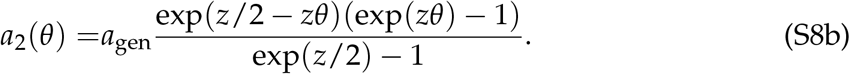

This is the trade-off parametrization for the four consumer traits *a*_1,*J*_, *a*_2,*J*_, *a*_1,*A*_, and *a*_2,*A*_ as depicted in figure 1 and it is used in all numerical calculations. We use *a*_gen_ = 0.001 throughout.

### S3 Invasion analysis

The evolutionary dynamics of the consumer traits in a population consisting of a single species - a monomorphic consumer population – can be studied using adaptive dynamics (Metz et al., 1992, 1996; Geritz et al., 1998). The starting point is to investigate the fate of a mutation ***θ***^m^ occurring in a resident population whose individuals are characterized by ***θ***^r^. Let us assume that mutations occur rarely such that the consumer population has reached its population dynamical equilibrium. At this equilibrium, the densities of the four resources are determined by the consumer’s feeding efficiencies and therefore its traits. We make this explicit in the notation by writing 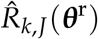 and 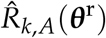. As long as the mutant sup-population is so small that it has a negligible effect on the densities of the four resources, its dynamics can be approximated by

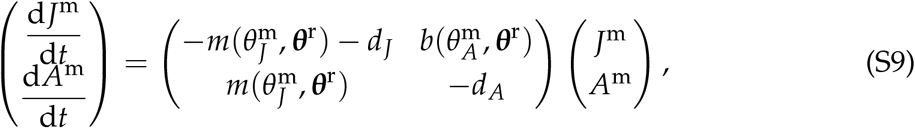

where

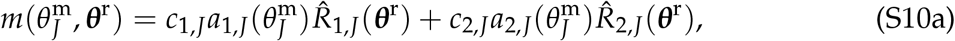

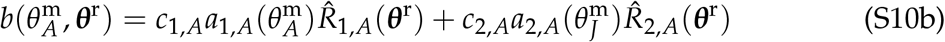

are the maturation and birth rate, respectively, of mutant individuals in an environment where the resource densities are set by the resident’s trait vector. Correspondingly,

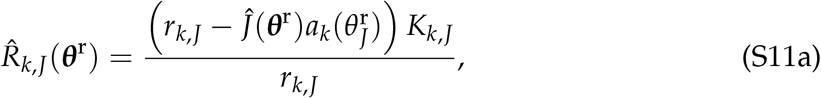

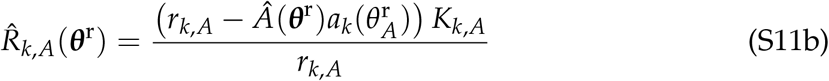

are the equilibrium densities of the four resource species where 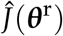 and 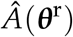 denote the equilibrium densities for juveniles and adults, respectively, of the resident population.

In the general case of a mutation appearing in a population consisting of *n* resident species the maturation and birth rate in equation (S9) read

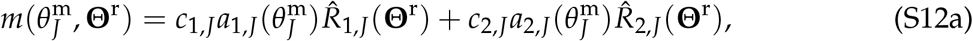

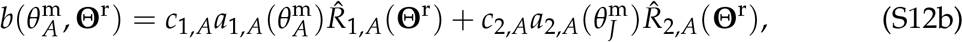

where 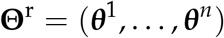 is the vector collecting the trait vectors of all *n* resident species. The expressions for the resource equilibria are obtained by replacing 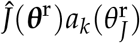 and 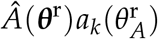 in equation (S11) with 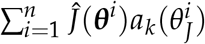 and 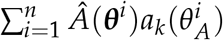, respectively, analogously to equation (S4).

The mutant sub-population has a positive probability to increase if the dominant eigenvalue of the matrix on the right-hand side of equation (S9), given by

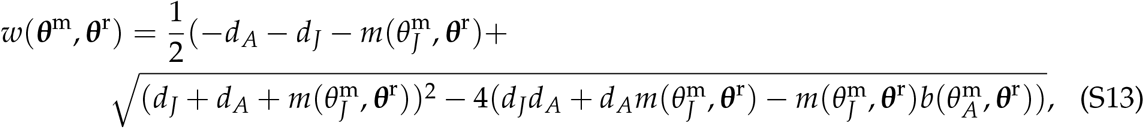

is positive and is doomed to extinction otherwise. We refer to *w*(***θ***^m^, ***θ***^r^) as invasion fitness. If *w*(***θ***^m^, ***θ***^r^) > 0 and the mutant trait vector ***θ***^m^ is sufficiently close to the resident trait vector ***θ***^r^, then a successfully invading mutant will ultimately replace the resident type (Dercole and Rinaldi, 2008), resulting in a trait substitution. Importantly, positive invasion fitness does not guarantee successful invasion since even a beneficial mutation can disappear due to demographic stochasticity as long as it is rare. This will be accounted for in our simulation algorithm (described in Appendix S5), where we incorporate a formula for the probability of successful invasion due to Saltini et al. (2022).

Repeated successful invasion of mutants and replacement of residents results in a trait substitution sequence. In the limit of rare mutations of very small effect occurring in very large resident populations this trait substitution sequence can be approximated by

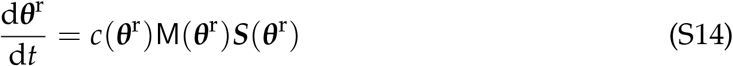

(Dieckmann and Law, 1996; Durinx et al., 2008). Here, *c*(***θ***^r^) is a real-valued function describing variation in the rate of mutations (e.g., due to variation in population size) and M(***θ***^r^) is the mutational variance-covariance matrix describing a symmetric distribution of mutations around the resident strategy. Finally, ***S***(***θ***^r^) denotes the fitness gradient – a column vector whose entries *S_J_*(***θ***^r^) and *S_A_*(***θ***^r^) are the partial derivatives of invasion fitness with respect to the mutant’s juvenile and adult trait values and evaluated at the resident trait vector, as per equations (6). These entries can be calculated as

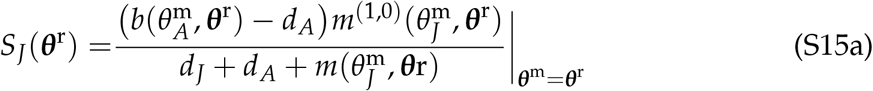

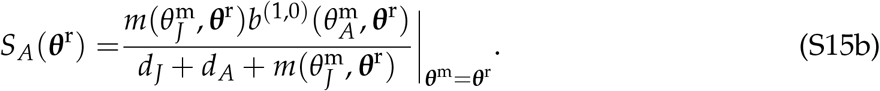

Here, the superscript (1,0) denotes the partial derivative with respect to the first argument. Note that 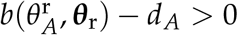 is a requirement for a consumer population not to go extinct. In the calculation of the fitness gradient we made use of the equality

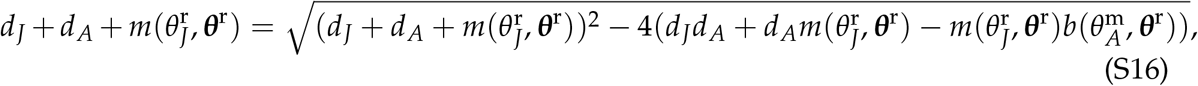

which follows from equation (S13) and noting that, by definition, *w*(***θ***^r^, ***θ***^r^) = 0. This equality is used throughout in the following analysis as it allows us to significantly simplify the expressions. The position of the singular points as shown in figure 2 and 4 are obtained by setting the corresponding component of the fitness gradient equal to zero and numerically solving for the trait value.

Equation (S14) describes an evolutionary dynamics that moves uphill on an ever changing fitness landscape. This dynamics comes to halt at trait vectors ***θ***\* for which the fitness gradient equals zero, ***S***(***θ***\*) = **0**. Such points are referred to as evolutionarily singular points. The classification of singular points in multi-dimensional trait spaces is in general complex (Leimar, 2009; Geritz et al., 2016; Vasconcelos and Rueffler, 2020). For our model, however, this task is straightforward as we show in the following. A singular point ***θ***\* is an attractor of the evolutionary dynamics, also called convergence stable, if it is an asymptotically stable fixed point of the evolutionary dynamics described by equation (S14) (Leimar, 2009). Convergence stability depends on the Jacobian matrix J of the fitness gradient with entries

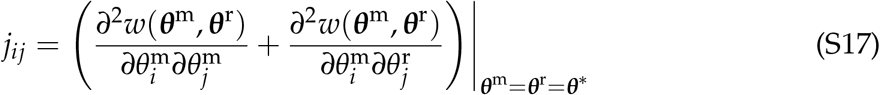

where *i, j* ∈ {*J, A*}. In particular, if the the symmetric part of J is negative definite, then a singular point ***θ***\* is an attractor of the canonical equation regardless of the mutational variance-covariance M (referred to as strong convergence stability; Leimar, 2009). Here, we show that for our model J is in fact a diagonal matrix such that negative definiteness amounts to both diagonal entries being negative. To see this, note that from the fitness gradient, given by equation (S15), it is clear that at a singular point 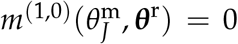 and 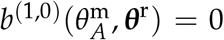 must hold. The two double-derivatives on the right-hand side of equation (S17) arise from differentiating the fitness gradient for a mutant and resident trait value, respectively. It is easy to see that the mixed derivatives (where we differentiate with respect to both a juvenile and an adult trait) contain either the derivative of the birth or maturation rate as a factor and that therefore *j_JA_* = 0 and *j_AJ_* = 0 must hold. Biologically, this means that the juvenile and adult traits are independent in their effect on fitness. For the diagonal entries of the Jacobian matrix we obtain

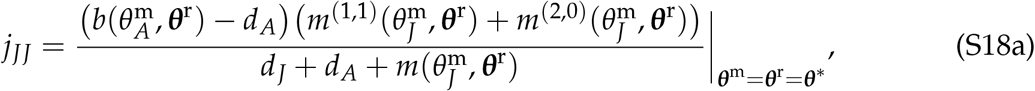

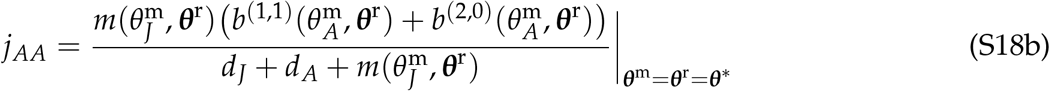

where 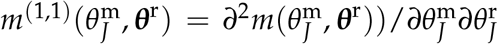 and 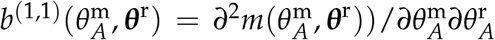. Furthermore, the superscript (2,0) denotes the second partial derivative with respect to the first argument. Thus, ***θ***\* is an attractor of the evolutionary dynamics independent of the mutational process if the diagonal entries of J are both negative, *j_JJ_* < 0 and *j_AA_* < 0. If at least one of the diagonal entries is positive, then ***θ***\* is an evolutionary repellor.

Inspecting the right-hand side of equations (S18a) and (S18b) allows for the following conclusion. Assuming that the abundance of a resource decreases the more specialized the consumer population becomes for that resource 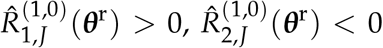, 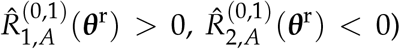, both 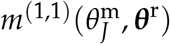 and 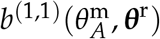 are negative. Since 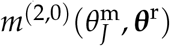 and 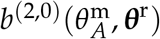 are negative for weak trade-offs and positive for strong trade-offs, it follows that the two diagonal entries of the Jacobian matrix are negative for weak trade-offs. In the next Appendix, we show that for fully symmetric parameter values the sign of *j_JJ_* and *j_AA_* at 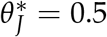 and 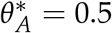 changes from positive to negative at some threshold value 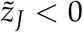 and 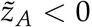, respectively.

Whether a singular point is invadable by nearby mutants is determined by the Hessian matrix H of invasion fitness which describes the local curvature of the fitness landscape around a singular point. The entries of H are

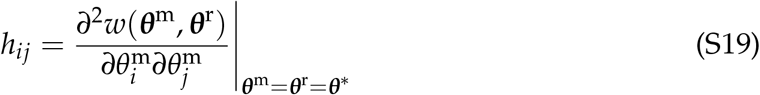

where *i, j* ∈ {*J, A*}. Using the same argument as for the off-diagonal entries of the Jacobian matrix, it is clear that *h_JA_* = 0 = *h_AJ_*. For the diagonal entries we obtain

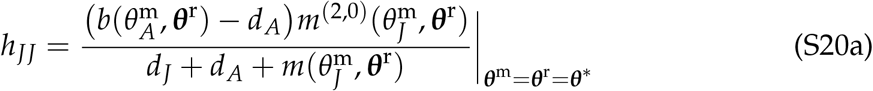

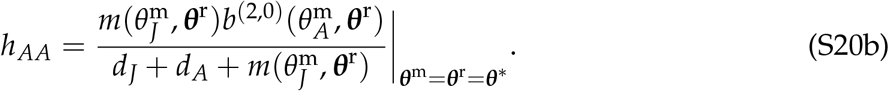

The sign of *h_JJ_* is determined by the sign of 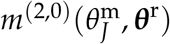, and the sign of *h_AA_* is determined by the sign of 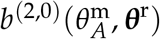. Since the second derivative of the feeding efficiencies *a*_1,*J*_(*θ*) and *a*_2,*J*_(*θ*) are positive for *z_J_* < 0 and negative for *z_J_* > 0, it follows that *h_JJ_* is negative if the trade-off is weak and positive if the trade-off is strong. The same argument holds for *h_AA_*. Thus, a singular point ***θ***\* is uninvadable if both trade-offs are weak and invadable if at least one of the trade-offs is strong. The evolutionary properties of the singular points shown in figure 2 and 4 are obtained by numerically evaluating the corresponding diagonal entry of the Jacobian and Hessian matrix.

### S4 Population size and evolutionary branching

The results in this appendix are derived under the symmetry assumptions that 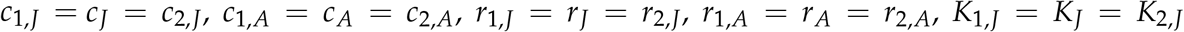, and 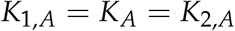. Due to this symmetry, and the symmetry that is inherent in the tradeoff parametrization presented in Appendix S2, the trait vector 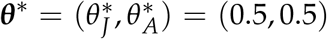 has to be singular point. The fact that a consumer population at this singular point is a perfect generalist for both resources at both life-stages induces further symmetry, which allows us to make the following definitions:

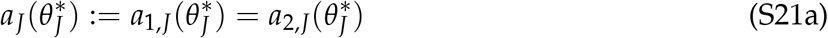

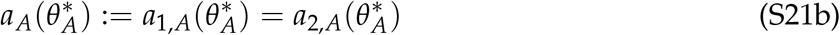

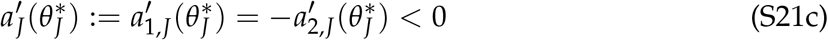

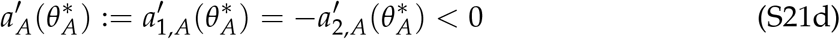

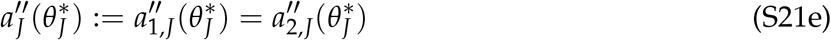

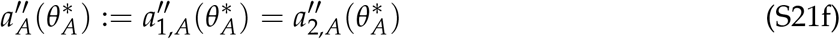

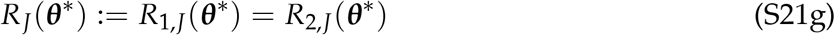

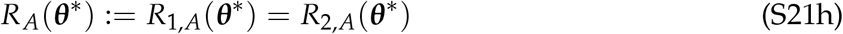

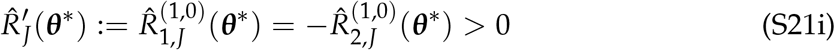

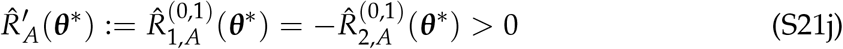

where ′ and ″ denote the first and second derivative, respectively, evaluated at ***θ***\*. The results concerning the trait values follow from the symmetry that is inherent in the trade-off parametrization while the results concerning the resource equilibria follow from the symmetry assumptions affecting equation (S11). The definitions in (S21) are used throughout this appendix.

We here take the perspective that only one of the traits evolves at a time while the other is fixed at the generalist trait value. This perspective allows us to compare the properties of the two singular points 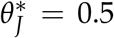 and 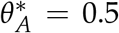. We first show that there exists a unique trade-off curvature 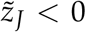 such that the entry of the Jacobian matrix *j_JJ_* (equation S18a), evaluated at 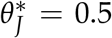, is negative for 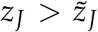 and positive for 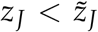. Since we have *h_JJ_* > 0 for *z_J_* < 0, it follows that the singular point 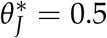 changes from an evolutionary branching point to an evolutionary repellor at 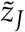. An analogous results holds true for the adult singular point 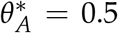. Second, we show that the interval of trade-off curvatures for which the generalist singular point is an evolutionary branching point is larger for the trait corresponding to the more abundant life-stage. In other words, 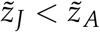 when 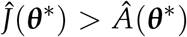 and 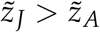 when 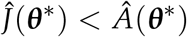.

Differentiating *j_JJ_* and *j_AA_* (equation S18) with respect to *z_J_* and *z_A_*, respectively, yields

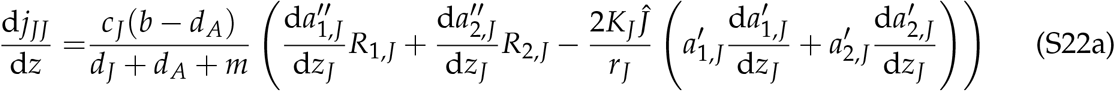

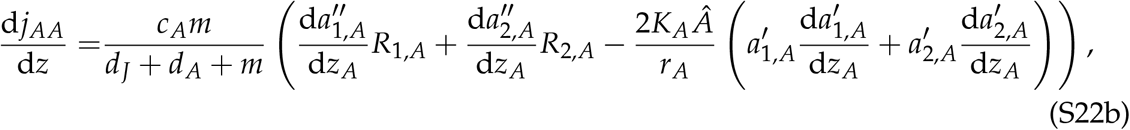

where all functions and derivatives are evaluated at 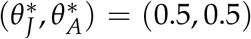. In simplifying these derivatives we made use of the fact that 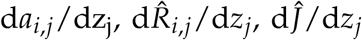 and 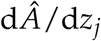 are all equal to zero, where *i* ∈ {1,2} and *j* ∈ {*J, A*}. This is due to our parametrization of the trade-off curve in which the value of *a_i,j_*(0.5) does not depend on the tradeoff curvature. Furthermore, 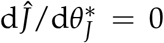 and 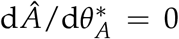 at the symmetric singular point. The right-hand side of equations (S22) is negative. This follows from 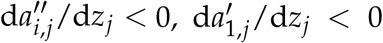 and 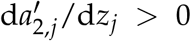. Thus, *j_JJ_* and *j_AA_* increase monotonically with decreasing values of *z_J_* and *z_A_*, respectively. Since *j_JJ_* < 0 and *j_AA_* < 0 for weak tradeoffs (*z_J_* > 0 and *z_A_* > 0), we can conclude that *j_JJ_* and *j_AA_* change sign at most once and – if they do – this occurs at a strong trade-off (*z_J_* < 0 and *z_A_* < 0).

Next, consider the trade-off curvature 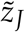 where at the singular point 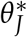 we have *j_JJ_* = 0. In other words, 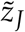 is the bifurcation point where the juvenile trait changes from convergence stable (*j_JJ_* < 0) to evolutionarily repelling (*j_JJ_* > 0). We will show that, for the same trade-off curvature in the adult trait, 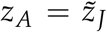, the singular point 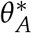 is evolutionarily repelling if the consumer population is juvenile dominated 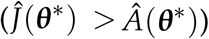 and convergence stable if the consumer population is adult dominated 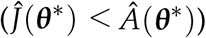. Since, according to the results from the previous section, these bifurcations occur for strong trade-offs where singular points are invadable, we obtain that the interval of trade-off curvatures resulting in an evolutionary branching point is larger in the trait corresponding to the dominant life-stage.

Given the equalities in (S21), it follows with equation (S18a) that *j_JJ_* equals zero if

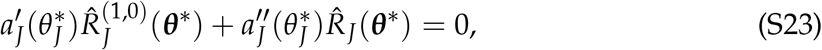

which is equivalent to

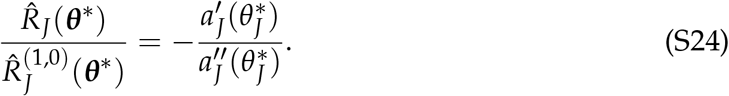

Similarly, it follows with equation (S18b) that *j_AA_* is negative if

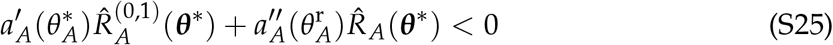

and positive if the reverse inequality is true. Thus, *j_AA_* < 0 is equivalent to

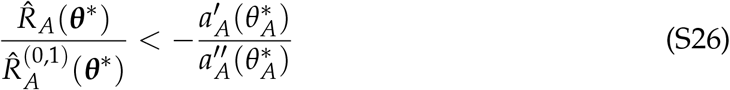

and if *j_AA_* > 0 the reverse inequality is true.

Since we use the same trade-off parameterization in the two life-stages and assume the same curvature, *z_J_* = *z_A_*, we have 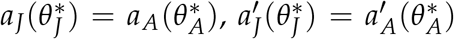 and 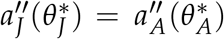. Thus, we can equate the right-hand side of equation (S24) with the right-hand side of inequality (S26) to obtain

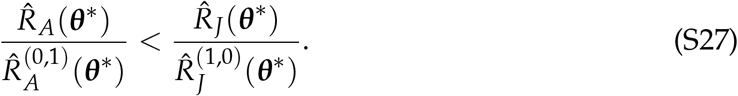

The interpretation of this inequality is as follows. For a given trade-off curvature 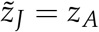 for which the the juvenile singular point is at the boundary between repelling and convergence stable (*j_JJ_* = 0) the adult singular point is convergence stable if inequality (S27) is true and repelling if the reverse inequality is true.

Next we insert the expressions for 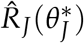 and 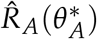 as given by equation (S11) and calculate the derivatives of these expressions where we can use the fact that 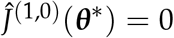 and 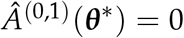 at the symmetric singular point. Inequality (S27) can then be rewritten as

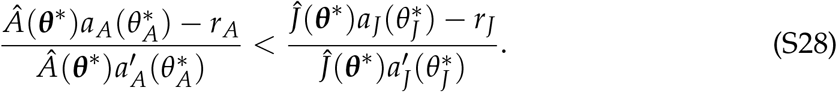

Using again the fact that we use the identical trade-off parametrizations and that 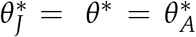 we have 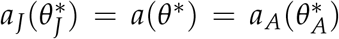 and 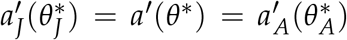. With this, inequality (S28) simplifies to

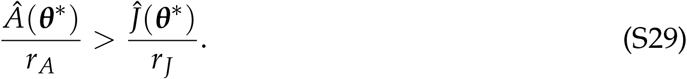

Thus, if inequality (S29) is true, then the adult singular point is convergence stable and, if it is reversed, then the adult singular point is evolutionarily repelling. We can conclude that with equal growth rates of the adult and juvenile resource (*r_J_* = *r_A_*) the trait corresponding to the more abundant life-stage has a larger interval of trade-off curvatures where it is convergence stable and, consequently, an evolutionary branching point.

### S5 Simulation algorithm

We use computer simulations performed in MATLAB (2021) to study the evolutionary dynamics in the two-dimensional trait space. Our simulations can be viewed as a stochastic implementation of equation (S14) where we allow for mutational steps of finite size. Simulations start with a monomorphic founding population consisting of 100 juvenile and 100 adult individuals with trait vector ***θ*** = (*θ_J_*, *θ_A_*) = (0.4,0.4). Then, we iteratively run the following simulation algorithm:

1. Population dynamical equilibrium: Numerically solve equation (S2) for its population dynamical equilibrium using Euler’s forward method with time increment *t* = 0.001. We consider the system to have reached equilibrium once the difference between the density of every species present in the population before and after an iteration of the population dynamics is less than 10^−5^.
2. Extinction: Remove any species *i* whose total population density (*J_i_* + *A_i_*) is less than the extinction threshold, set to 0.01.
3. Determine the parent species *j* giving rise to the next mutant: The parent species producing the next mutant is drawn randomly with probabilities proportional to the number of offspring produced by each species. The index *j* ∈ {1,…,*n*} is given by the lowest integer to satisfy 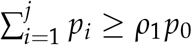. Here, 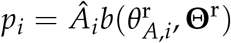 with the function *b* given by equation (S12b) and resource densities as obtained at the end of step 1. Thus, *p_i_* is the expected number of offspring by adults of species *i* at population dynamical equilibrium. Furthermore, *ρ*_1_ is a uniformly distributed random number between 0 and 1, and 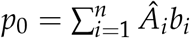.
4. Determine potential mutant phenotypes: Construct the four trait vectors 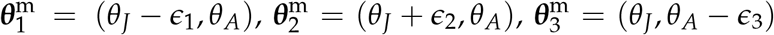 and 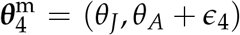, where *ϵ*_1_,…,*ϵ*_4_ are drawn from a Gaussian distribution with the parent trait value as mean and variance 0.02.
5. Assign mutant establishment probabilities: Each of the four mutants 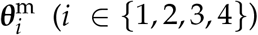 determined in the previous step has a certain probability to escape demographic stochasticity while rare and successfully establish in the population. The expression for this establishment probability is due to Saltini et al. (2022) and given by

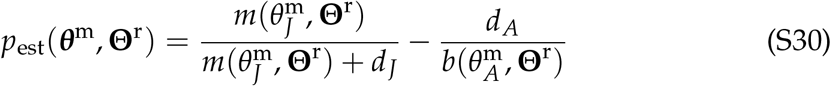

with the functions *m* and *b* given by equation (S12).
6. Determine the next mutant: The next mutant appearing in the population is drawn from the four candidate mutants with probabilities proportional to their establishment probabilities. The index *k* is the lowest integer to satisfy 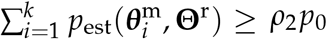. Here, *ρ*_2_ is a uniformly distributed random number between 0 and 1, and 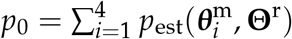.
7. Add mutant: Add one juvenile individual with the mutant phenotype vector to the consumer population.
8. Go back to step 1.

Each simulation of the symmetric case runs for 500 mutation events, and each of the asymmetric case runs for 700 mutation events.

